# Culture of human nasal olfactory stem cells and their extracellular vesicles as advanced therapy medicinal products

**DOI:** 10.1101/2022.08.05.502926

**Authors:** Charlotte Jaloux, Maxime Bonnet, Marie Vogtensperger, Marie Witters, Julie Veran, Laurent Giraudo, Florence Sabatier, Justin Michel, Romaric Lacroix, Corinne Chareyre, Regis Legré, Gaelle Guiraudie-Capraz, François Féron

**Affiliations:** Aix Marseille Univ, CNRS, INP, UMR 7051, Institut de Neuropathophysiologie, Equipe Nasal Olfactory Stemness and Epigenesis (NOSE), CEDEX 15, F-13344 Marseille, France; Department of hand surgery and reconstructive surgery of the limbs - La Timone University Hospital - Assistance Publique Hôpitaux de Marseille, 264, rue Saint-Pierre, 13005 Marseille, France; Aix Marseille Univ, CNRS, ISM, UMR 7287, Institut des Sciences du Mouvement : Etienne-Jules MAREY, Equipe Plasticité des Systèmes Nerveux et Musculaire (PSNM), Parc Scientifique et Technologique de Luminy, Faculté des Sciences du Sport de Marseille, CEDEX 09, F-13288 Marseille, France; Cell Therapy Department, Hôpital de la Conception, AP-HM, INSERM CIC BT 1409, 147 Bd Baille, 13005, Marseille, France; Department of Hematology and Vascular Biology, CHU la Conception, APHM, Marseille, France; Aix-Marseille Université, C2VN, UMR-1263, INSERM, INRA 1260, UFR de Pharmacie, Marseille, France; Aix Marseille University, Assistance Publique des Hôpitaux de Marseille, Institut Universitaire des Systèmes Thermiques Industriels, Department of Otorhinolaryngology and Head and Neck Surgery, La Conception University Hospital, Marseille, France

## Abstract

The olfactory ecto-mesenchymal stem cell (OE-MSC) are mesenchymal stem cells originating from the lamina propria of the nasal mucosa. They have neurogenic and immune-modulatory properties and showed therapeutic potential in animal models of spinal cord trauma, hearing loss, Parkinsons’s disease, amnesia, and peripheral nerve injury.

In this paper we designed a protocol that meet the requirements set by human health agencies to manufacture these stem cells for clinical applications.

Once purified, OE-MSCs can be used *per se* or expanded in order to get the extracellular vesicles (EV) they secrete. A protocol for the extraction of these vesicles was validated and the EV from the OE-MSC were functionally tested on an *in vitro* model.

Nasal mucosa biopsies from three donors were used to validate the manufacturing process of clinical grade OE-MSC. All stages were performed by expert staff of the cell therapy laboratory according to aseptic handling manipulations, requiring grade A laminar airflow. Enzymatic digestion provides more rapidly a high number of cells and is less likely to be contaminated. Foetal calf serum was replaced with human platelet lysate and allowed stronger cell proliferation, with the optimal percentage of platelet lysate being 10%. Cultivated OE-MSCs are sterile, highly proliferative (percentage of CFU-F progenitors was 15,5%) and their maintenance does not induce chromosomal rearrangement (karyotyping and chromosomal microarray analysis were normal). These cells express the usual phenotypic markers of OE-MSC. Purification of the EVs was performed with ultracentrifugation and size exclusion chromatography. Purified vesicles expressed the recognized markers of EVs (Minimal Information for Studies of Extracellular Vesicles (“MISEV”) guidelines) and promoted cell differentiation and neurite elongation in a model of neuroblastoma Neuro2a cell line.

We developed a safer and more efficient manufacturing process for clinical-grade olfactory stem cells, these cells can now be used in humans. A phase I clinical trial will begin soon.An efficient protocol for the purification of the OE-MSC EVs have been validated. These EVs exert neurogenic properties *in vitro*. More studies are needed to understand the exact mechanisms of action of these EVs and prove their efficacy and safety in animal models.

## 1. Introduction

Applied in the very first clinical trial based on cell therapy, performed in the mid-1950s (1), mesenchymal stem cells (MSC) opened a new field of therapeutic interventions. Since then, hundreds of MSC-based transplantations have been performed, in a wide range of diseases or trauma (2). And, while grafting experiments flourished, the family of MSC never ceased to expand. Initially restricted to bone marrow, mesenchymal stem cells were identified in numerous tissues, including umbilical cord, amniotic fluid, placenta, muscle, liver, skin, face and many others (3).

A subtype of mesenchymal stem cells, originating from the neural crest and mostly located in the face, has been isolated and categorized under the name of ecto-mesenchymal stem cells (4). These cells, previously named mesenchymal–neural precursors, can differentiate into ectoderm and mesoderm cell types (5) and, for clinicians willing to repair the nervous system, one specific subtype – the olfactory ecto-mesenchymal stem cell (OE-MSC) – is of particular interest. OE-MSCs belong to a nervous tissue and are involved in the permanent neurogenesis that takes place all lifelong in the olfactory mucosa (6). Located in the nasal cavity, this tissue is easily biopsied in vigil individuals, under local anesthesia (7).

By the end of the last century, multipotent progenitors able to give rise to neural and non-neural cells were isolated from rodent (8) and human (9,10) olfactory mucosa. Later on, these progenitors were characterized as a member of the ecto-mesenchymal stem cell family with neurogenic (11) and immune-modulatory (12) properties. While the phenotype of OE-MSCs was extensively characterized, their therapeutic potential was assessed in animal models of spinal cord trauma (13,14), hearing loss (for a review, (15)), Parkinsons’s disease (16), amnesia (17) and peripheral nerve injury (18–22).

In anticipation of clinical applications, the time has come to design a protocol that will meet the requirements set by human health agencies. Previously, our team described methods for cultivating human OE-MSCs (7,23). We also demonstrated *in vitro* how they can cross the blood-brain barrier (24) and be transdifferentiated into dopaminergic neurons (25). However, the manufacture of such a medicinal product must strictly follow good manufacturing practices (GMP) from its development through to the marketing authorization. According to the European Directive No. 1394/2007, stem cell is considered as an advanced therapy medicinal product (ATMP) and as such must comply with the rules and guidelines of the European Medicines Agency. We thus devised an innovative protocol with new methods and compounds that guarantee the quality and safety of OE-MSCs and their extracellular vesicles.

Once purified, OE-MSCs can be used *per se* or expanded in order to get the extracellular vesicles (EV) they secrete. Discovered in the mid-20^th^ century (26), EVs are known for carrying bioactive molecules - proteins, lipids, metabolites, nucleic acids - that act as signalling mediators in various metabolic pathways (27). Within the central nervous system, EVs are involved in synaptic plasticity, myelination, neurogenesis and neuroinflammation (28). Extracellular vesicles from mesenchymal stem cells promote peripheral nerve regeneration, angiogenesis, myelination (29–31) and modulate inflammation (32,33) *via* the release of trophic and immunomodulatory factors.

With regard to the olfactory stem cells, it has been observed that their EVs are CD63^+^, CD9^+^, calnexin^-^ and CD81^-^ (34–36). They carry at least 304 proteins, among which plasminogen activator inhibitor 1 (SERPINE 1), insulin-like growth factor binding protein family members (IGFBP4,5), epidermal growth factor receptor (EGFR), neurogenic locus notch homolog protein 2 (NOTCH 2), apolipoprotein E (APOE) and heat shock protein HSP90-beta (HSP90AB1) play a key role in angiogenesis, cell proliferation and differentiation, apoptosis and inflammation (36). *In vitro*, olfactory stem cells-derived EVs increase the proliferation and migration of human cerebral microvascular endothelial cells (36). When transplanted in mouse models of inflammatory colitis or Sjogren’s syndrome, they act as antiinflammatory agents and slow down the progression of the symptoms (34,35). To move a step further towards brain and nerve repair, we assessed *in vitro* the differentiating potential of OE-MSC EVs. For that purpose, we used N2a (37), a fast-growing neuroblastoma cell line. When differentiated, these cells display many properties of neurons, including neurofilaments. Extracellular vesicles were loaded in the culture medium and neurite differentiation was monitored, at various time points. In parallel, a similar experiment was performed with the conditioned medium of OE-MSCs.

## 2. Materials and methods

### 2.1 Study Design and Donor qualification

Initiated in December 2018, a prospective study entitled NOSE (ClinicalTrials.gov Identifier: NCT04020367), aimed to validate the manufacture of olfactory stem cells, using good manufacturing practices (38), in order to use them as an Advanced Therapeutic Medicinal Product (ATMP) for repairing peripheral nerves.

Five healthy adult individuals undergoing a turbinoplasty or a septoplasty under general anesthesia were included. A complete written and oral information on the goal and procedure of this research was provided to the participants and a signed informed consent was obtained from all of them, prior to their involvement in the study. All procedures were approved by the ethical committee (Comité de Protection des Personnes, file 2018-A00796-49) and the national competent authorities. For each individual, an olfactory mucosa biopsy and a blood sample (for genetic studies) were harvested.

Exclusion criteria were: central nervous system pathologies, pregnancy or breastfeeding women, chronic rhinosinusitis or acute rhinosinusal infection, anti-thrombotic treatment, allergy to amide-type local anesthetics or to one of their excipients, previous surgical treatment on ethmoid and/or medium turbinates, patients with porphyria or uncontrolled epilepsia (contraindications to xylocaine and naphazoline applications), previous cervical or encephalic radiotherapy, positivity to viral serologies (VIH 1, VIH 2, VHB, VHC, HTLV1) or hemostasis anomalies (on blood works).

All manufacturing processes were achieved in the LCTC (Laboratory of Culture and Cell Therapy), which is the cell therapy department of La Conception University Hospital, this laboratory is approved by the health authorities (ANSM = French national agency for the safety of medicines) to produce ATPMs (FR01303M for ATMP “hospital exemption”, FR 01304 M for investigational ATMP).

### 2.2 Collection of olfactory mucosa biopsies

Biopsies were performed unilaterally by an ENT surgeon after informed consent (Nose study), at the level of the middle turbinate arch, using a Morscupula forceps and an endoscope, as described before (7). When the patient was deeply anesthetized and just before the planned procedure, a 2 mm^2^ biopsy was collected and transferred in a sterile tube filled with culture medium + penicillin + gentamycin (#, alpha MEM Macopharma BC0110020). The tube was then transferred to the Laboratory of Culture and Cell Therapy (LCTC, AP-HM, using a transport container for processing (#, ISOS 03).

### 2.3 Olfactory ecto-mesenchymal stem cells

#### 2.3.1 Olfactory ecto-mesenchymal stem cells manufacturing process

The whole culture of olfactory stem cells was performed in aseptic conditions to further therapeutic applications. After biopsy excision, all stages were performed by expert staff of the LCTC according to aseptic handling manipulations, requiring grade A laminar airflow. All consumables and raw materials were marketed for therapeutic use by suppliers

Our previously published protocols (7,23,24) were modified in order to 1) use agreed raw materials and processing raw materials and cells bank according to ATMP good manufacturing practices (38) and 2) speed up the proliferation of stem cells. With these intentions in mind, we compared two modes of cell individualization (enzymatic dissociation of the mucosa *versus* explant culture under coverslip, figure 1) and two cell proliferation-inducing agent (therapeutic serum *versus* platelet lysate). Enzymatic dissociation was performed using 1 mL of collagenase NB6 (1U/mL, Nordmark Biochemicals). After a one-hour incubation at 37 °C, the tissue was mechanically dissociated, and the enzymatic activity was stopped by adding 9 mL of Ca-free and Mg-free PBS. The cell suspension was centrifuged at 400g for 5 min and the pellet was resuspended in a culture medium, supplemented with penicillin + gentamycin and either serum (Invitrogen) or platelet lysate (Macopharma®, Tourcoing, France), at various concentrations. The following culture media were tested: Gibco Stem Pro (#A1014201) (medium 1), Gibco Stem Pro + CellStart (#A1014201) (medium2), DMEM/HAM F12 with 10% SVF (10094-142) with or without FGF2 (Miltenyi 130-093-838), αMEM (BC0110020) with platelet lysate. Two lines of platelet lysate (PL30 = BC0190020 vs PL100 = BC0190030), with or without gamma ray irradiation, were compared.

**Figure 1.**
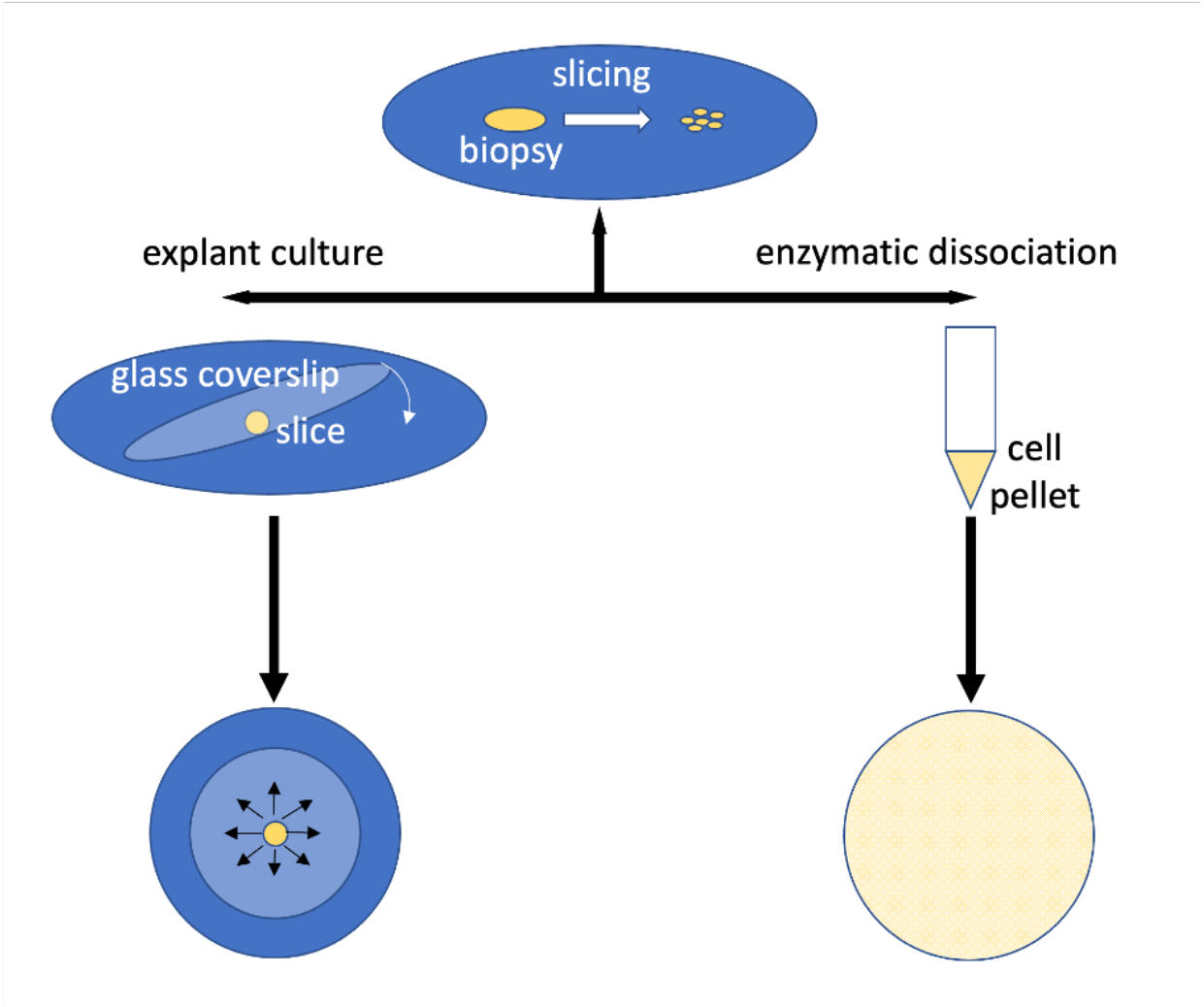
Schematic description of the two procedures tested for the isolation of olfactory stem cells. Biopsies were chopped and slices were either cultivated under glass coverslips or enzymatically dissociated.

For comparison purpose, two similar biopsies, collected at the same location within the nasal cavity of the same donor, were cultivated in parallel, according to the two methods described above. Inter-donor reproducibility was assessed by using nasal biopsies of 2 or 3 donors for each culture.

Proliferation rate was assessed by measuring the doubling time at Day 5 post-plating.

#### 2.3.2 Quality controls of cultivated olfactory ecto-mesenchymal stem cells

According to the recommendations of the European medicine agency, the quality of the ATMP must be evaluated according to several criteria. Quality criteria were assessed here following the subsequent methods.

##### Cell number and viability

Both these parameters were assessed with the LUNA-FL™ Automated Fluorescence Cell Counter, dual fluorescence counting was achieved with Acridine Orange/Propidium Iodide Stain.

##### Cell phenotyping

Characterization of OE-MSCs was performed by flow cytometry. Aliquots of 5 × 10^5^ OE-MSCs per tube were suspended in 100 μL of PBS and stained for 20 min at room temperature in the dark with the DRAQ5 nuclear marker and pre-prepared antibodies or corresponding isotype controls in matched concentrations. The primary antibodies were anti-CD31 (Réf. IM1431U / Fournisseur Beckman Coulter / Clone 5.6E), anti-CD34 (IM2709U / Beckman Coulter / 581), anti-CD45 (A07785 / Beckman Coulter / J33), anti-CD90 (IM1839U / Beckman Coulter / F15-42-1-5), anti-CD146 (A07483 / Beckman Coulter / TEA1-34), anti-S100A4 (ab41532 / Abcam / polyclonal) and anti-nestin (Nestin humain - clone 10C2 - Réf. MAB5326 – 1mg/ml). They were conjugated with the following fluorochromes: fluorescein isothiocyanate (FITC), phycoerythrin (PE), phycoerythrin-Texas Red-X (ECD), and phycoerythrin-Cyanin 5.1 (PC5). Flow cytometry was performed on a NAVIOS instrument (Beckman Coulter, Brea, CA, USA) and data files were analyzed using Kaluza software (Beckman Coulter) with a multiparameter gating strategy.

##### Sterility and microbial assays

The following sterility and microbial assays were performed using the supernatant of each OE-MSC culture:

- Aerobic and anaerobic blood cultures, assessing the presence of germs with the BACT/ALERT® 3D kit (Biomérieux)
- Turbidimetry assay determining the presence of endotoxins
- PCR assay for Mycoplasma sp. detection

##### Clonogenicity assay

The clonogenic potential of OE-MSCs was assessed using the colony formation assay with CFU-F. OE-MSCs manufactured with the irradiated PL100 were plated/seeded at a concentration of 100 cells/well in 6 well plates. The culture medium was DMEM/Ham’s F12 with 10% fetal calf serum.

Incubation was performed at 37°C under 5% CO2. The medium was renewed between Day 3 and Day 5 and then every 2 to 3 days, for 14 days. At the end of the culture, cells were Giemsa-stained and colonies with at least 50 cells were counted with an inversed-microscope. CFU-F percentage was determined for each well.

##### Genetic stability of OE-MSCs

OE-MSCs manufactured with the irradiated PL100 were evaluated for genetic stability using two cytogenetic assays: conventional karyotype and Array-CGH (comparative genomic hybridization).

### 2.4 Olfactory ecto-mesenchymal stem cells derived extracellular vesicles

#### 2.4.1 Purification OE-MSC-derived EVs

The procedure is summarized on figure 2. For 4 days, OE-MSCs from 3 donors were cultivated in αMEM, supplemented with platelet lysate (5%), heparin (2UI/ml), gentamycin and penicillin. The next four days, the platelet lysate was replaced by a supplementation with ITS (insulin, transferrin, selenium). Supernatants were collected and centrifuged at room temperature for 5 min at 300 g to remove cells and then centrifuged again for 15 min at 2,500 g to remove cell debris. Subsequently, supernatants were centrifuged for 90 min at 100,000 g at 4 °C and EV pellets were resuspended in PBS. EV samples were subjected to size-exclusion chromatograhy using qEV original size exclusion columns (Izon, Cambridge, MA, USA) for the removal of soluble proteins. Then, 500 μl fractions were successively collected after the addition of EV samples on the top of the columns. Fractions 7 and 9 enriched in EVs were collected in a final volume of 1,5 mL.

**Figure 2.**
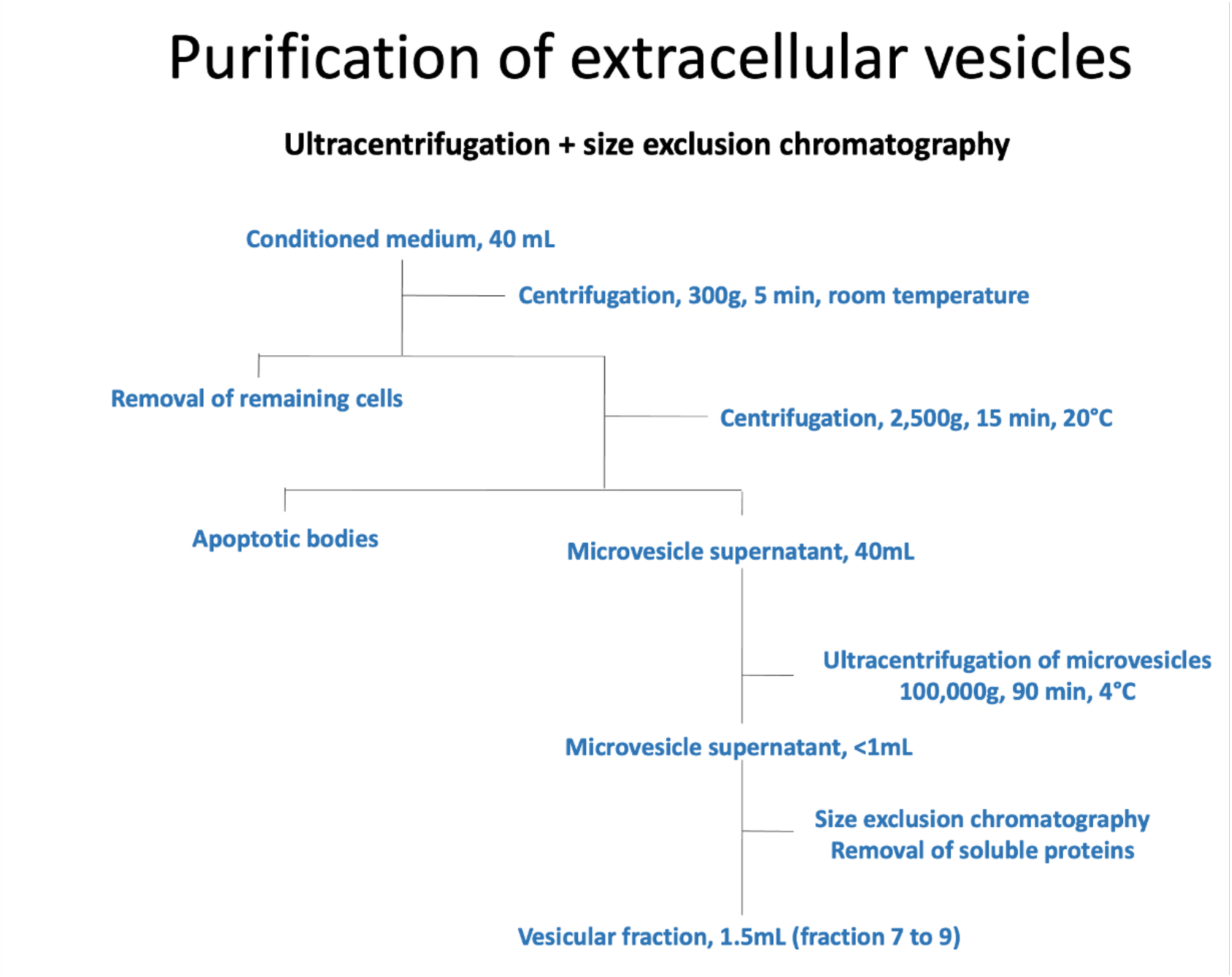
Step by step procedure for isolating extracellular vesicles.

#### 2.4.2 Characterization of OE-MSC EVs

We have submitted all relevant data of our experiments to the EV-TRACK knowledgebase (EV-TRACK ID: EV210203) (Van Deun J, et al. EV-TRACK: transparent reporting and centralizing knowledge in extracellular vesicle research. Nature methods. 2017;14(3):228-32 (39)).

##### Size determination

Size clustering of OE-MSC EVs was performed using the qNano instrument (Izon Science Ltd, Christchurch, New Zealand) and the tunable resistive pulse sensing (TRPS) (40). We followed manufacturers’ recommendations and NP400 nanopore membranes stretched between 43–47 nm were used. Voltage was set in 0.3–0.5 V to achieve a stable current 90–120 nA and the pressure at 0.8 kPa, with the root mean square noise below 10 pA. Calibration beads used were CPC400 (mean diameters 350 nm). Running electrolyte was PBS filtered under 0.1 μm. Measurement and analysis were performed with Izon Control Suite V3.3.3.2001 Software.

##### Flow cytometry

OE-MSC EVs and OE-MSCs were analyzed using high-sensitivity flow cytometry as previously described (41). Thirty microliters of EV samples were incubated with the appropriate amount of specific antibodies and 10 μL of FITC-bound Annexin V reagent (Tau Technologies, Netherlands). Each stained sample was analyzed on a 3-laser Navios flow cytometer (Beckman-Coulter), based on a protocol standardized with Megamix-Plus FSC beads (BioCytex, Marseille, France). The following fluorescently-tagged antibodies were used: PE-tagged anti-CD59, PE-tagged anti-CD29, APC-tagged anti-CD41 (Beckman Coulter).

##### Western blotting

Purified EVs were lysed with RIPA buffer, separated on 4-12% NuPAGE 1.5 × 10 wells gel (Novex), under reducing or non-reducing conditions and transferred onto the nitrocellulose membrane (0.45 μm, Amersham). After a blocking step, membranes were incubated overnight at 4°C with antibodies for the designated antigens INTEGRIN α5 (1:1000, #4705), INTEGRIN β3 (1:1000, #4702), β-ACTIN(1:1000, #8457), β-TUBULIN (1:1000, #2146) purchased from Cell Signaling Technology, GAPDH (1:1000, #MAT11114), ALBUMIN (1:1000, #MAT532531), CD63 (1:1000, #MAT10628D), CD81 (1:1000, #MAT10630D), GAPDH (1:1000, #MAT11114) purchased from ThermoFisher, CD9 (1:1000, #SC59140) purchased from CliniScience and CAVEOLIN1 (1:1000, #03600) purchased from Invitrogen. Next, the horse radish paroxidase (HRP)-conjugated secondary antibody (1:3000-1:5000, Pierce, ThermoFisher) was added for 1h at room temperature. Immunocomplexe were visualized by enhanced chemuluminescence (ECL or ECL femto) according to the manufacturer’s instruction (Pierce, Rockford, IL, USA). Specific bands were detected using a G-Box Imaging System (GeneSys, Cambridge, UK).

#### 2.4.3 Neuronal differentiation assays

We performed *in-vitro* experiments with the mouse neuroblastoma Neuro2a cell line to assess the neurotrophic effects of OE-MSC extracellular vesicles.

##### Culture of N2a cell line

Neuro 2A (N2a) is a neural crest-derived murine cell line that is used for studying neuronal differentiation, axonal growth and neural signaling pathways. N2a were grown and maintained in a humidified incubator in 75 cm^2^ flasks with Dulbeco’s modified eagle’s medium (DMEM)-Glutamax supplemented with 10% fetal calf serum, 1% sodium pyruvate (100 mM), 1% penicillin (100 U/ml) and streptomycin (100 μg/ml). Passaging was performed with trypsin/EDTA.

##### N2a plating

N2a cells were plated in 12-well culture plates at the concentration of 20,000 cells/3.6 cm^2^ well in DMEM including glucose, 200 μM L-glutamine, 5% fetal calf serum, penicillin (100 U/ml) and streptomycin (100 μg/ml). The medium was changed the following day to set up a new culture medium. These conditions were maintained for 4 days with a renewal of the medium on the second day if necessary. Drug-induced differentiation was performed by adding 2 mM dibutyryl-cAMP to the culture medium.

##### Vesicle-induced differentiation

OE-MSCs from 3 donors were cultured in αMEM supplemented with 5% platelet lysate and heparin (20 IU/ml). On day 4, cells were cultured in αMEM supplemented with 1% ITS (insulin, transferrin, selenium). After 96h of culture, supernatants were collected and purified as previously described (2.6). For the differentiation assay, 3 experimental conditions were performed in triplicate. For this purpose, a mixture of culture medium containing 1% SVF and supplemented with 20,000, 200,000 and 2,000,000 EVs respectively were inserted into the wells containing N2a cells.

##### Neurite length assay

At Day 2, Day 3 and Day 4, N2a cells were photographed with a phase contrast light microscope with a ×10 or x20 magnification objective. (Leica DMI4000B microscope, Leica DFC340 FX digital camera). Five randomly selected fields per well were analyzed in a double-blind manner, using the Image J software and the cell counter plugin. Only cells with a neurite extending over twice the size of the diameter of the cell body were taken into consideration. Neurite length was measured with the “NeuronJ” plugin of Image J software. Additional measurements (total number of extensions per field, number of extensions per cell) were performed on a random sample representative of all conditions under study.

## 3 Results

### 3.1 A safe and high efficiency production of olfactory ecto-mesenchymal stem cells

#### 3.1.1 Enzymatic digestion provides more rapidly a high number of cells

As reported in the table 1 below, the explant technique takes more time to reach confluency after the initial plating and, as a consequence, a longer delay is required to get over 5 million cells. We also compared two enzymes for passaging – trypsin/EDTA *versus* TrypLE – and found no significant difference between the two cocktails: in both cases, a full dissociation was achieved, the viability was over 99% and the number of cells at Day 4 reached similar values. These findings were replicated with OE-MSCs originating from three different donors.

**Table 1.** Comparison of Enzymatic digestion and Explant culture conditions for the proliferation of the olfactory ecto-mesenchymal stem cell.

|  | Enzymatic digestion | Explant culture |
| --- | --- | --- |
| Time before passage | 14 days | 18 days |
| Time at confluency after passage | 21 days | 24 days |
| Total number of cells | 7,8 10<sup>6</sup> | 5,6 10<sup>6</sup> |

#### 3.1.2 Human platelet lysate as a potent alternative to fetal calf serum

When compared with serum, platelet lysate (v/v,10%) induces a stronger cell proliferation: for the two platelet lysates, the fold change is 3.1 and 4.1, respectively (Figure 3A). Platelet lysate PL30 (PL1 in figure 3) was less effective than platelet lysate PL100 (PL2 in figure 3). Even when serum is complemented with FGF2, it remains less favorable than platelet lysate. Increasing the percentage of platelet lysate, up to 20%, is not beneficial since, in this condition, the doubling time augments (data not shown). Conversely, reducing the percentage of platelet lysate to 7.5% has no detrimental effect on cell proliferation (Figure 3B). An inter-batch variability is noticeable, but it tends to diminish at the higher concentrations (Figure 3C). Gamma ray irradiation, a procedure in line with hospital health and safety guidelines, was also tested. As shown on Figure 3D, cell proliferation remains almost steady after sterilization of the lysate.

**Figure 3.**
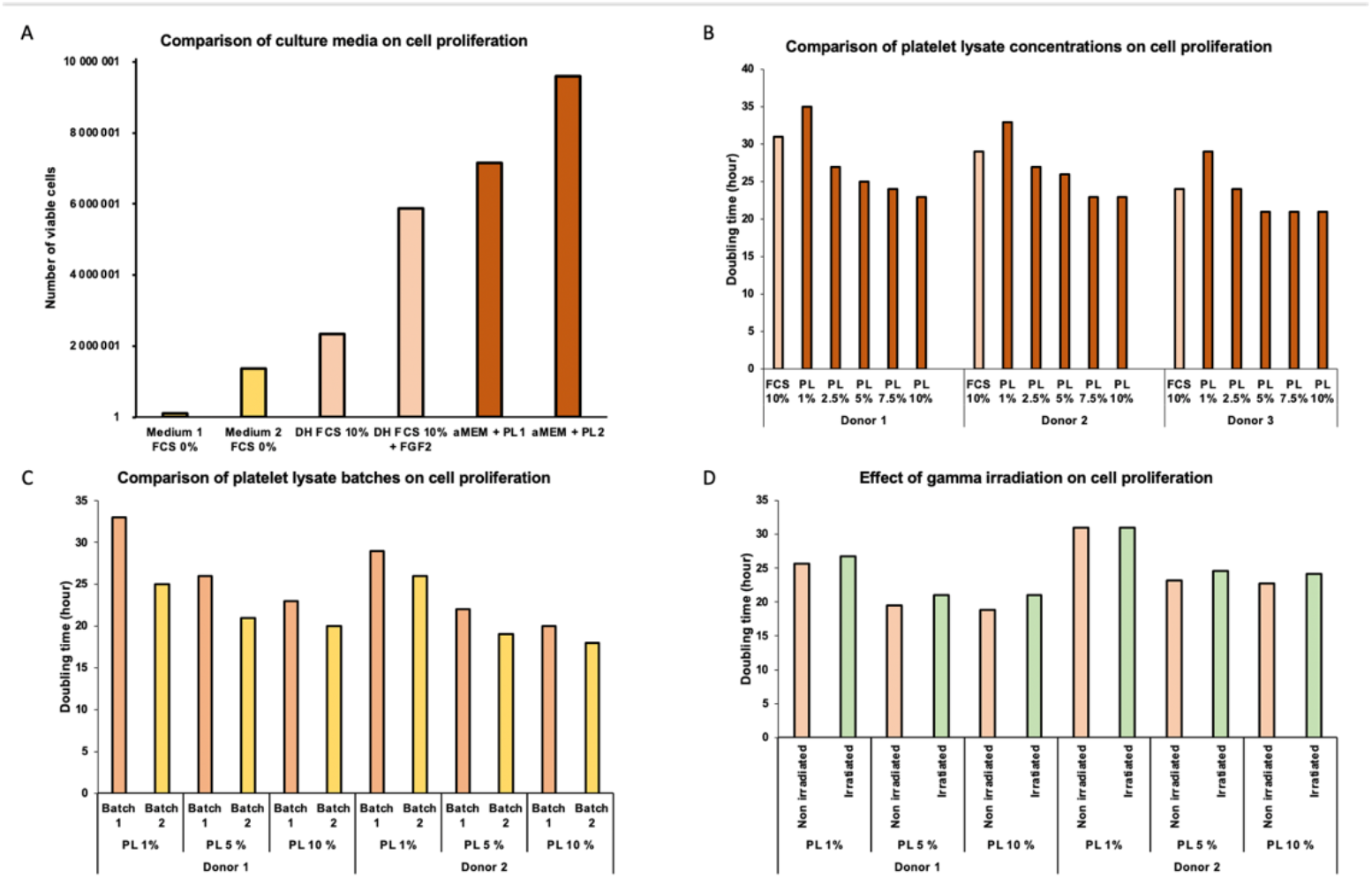
Comparison of culture conditions on cell proliferation. A) Effect of culture medium supplemented with platelet lysate on cell proliferation was compared with standard media including 10% fetal calf serum. B) Comparison of five concentrations of platelet lysates on cell proliferation. Doubling time is reduced when platelet lysate is equal to or above 5%. C) Inter-batch variability on cell proliferation. D) Effect of the sterilization procedure on cell proliferation. Gamma irradiation has no detrimental effect on doubling time.

#### 3.1.3 Cultivated OE-MSCs are sterile, highly proliferative and their maintenance does not induce chromosomal rearrangement

Sterility and microbial assays ruled out the presence of Mycoplasma sp., endotoxins, bacteria, and fungi. All final cultures were sterile. OE-MSCs display a high self-renewal activity: the mean percentage of CFU-F progenitors among the originally seeded cells was 15,5%. Putative genetic anomalies were assessed by using the techniques of karyotyping and chromosomal microarray analysis. None of these techniques detected polyploidy and/or deletions or duplications, at passage 3.

#### 3.1.4 Purified cells express the recognized markers of OE-MSCs

As expected, OE-MSCs are negative for CD31, CD34 and CD45, recognized markers of hematopoietic stem cells, and positive for CD 90, CD146 and CDx, surface markers of olfactory stem cells (Figure 4). In addition, OE-MSCs express the protein S100A4, known as an intracellular marker of this specific cell type. However, to make sure that the culture was not contaminated by fibroblasts, a cell type present in the *lamina propria* that produce the proteins described above, we assessed the expression of NESTIN. It was observed that only OE-MSCs express this stem cell marker.

**Figure 4.**
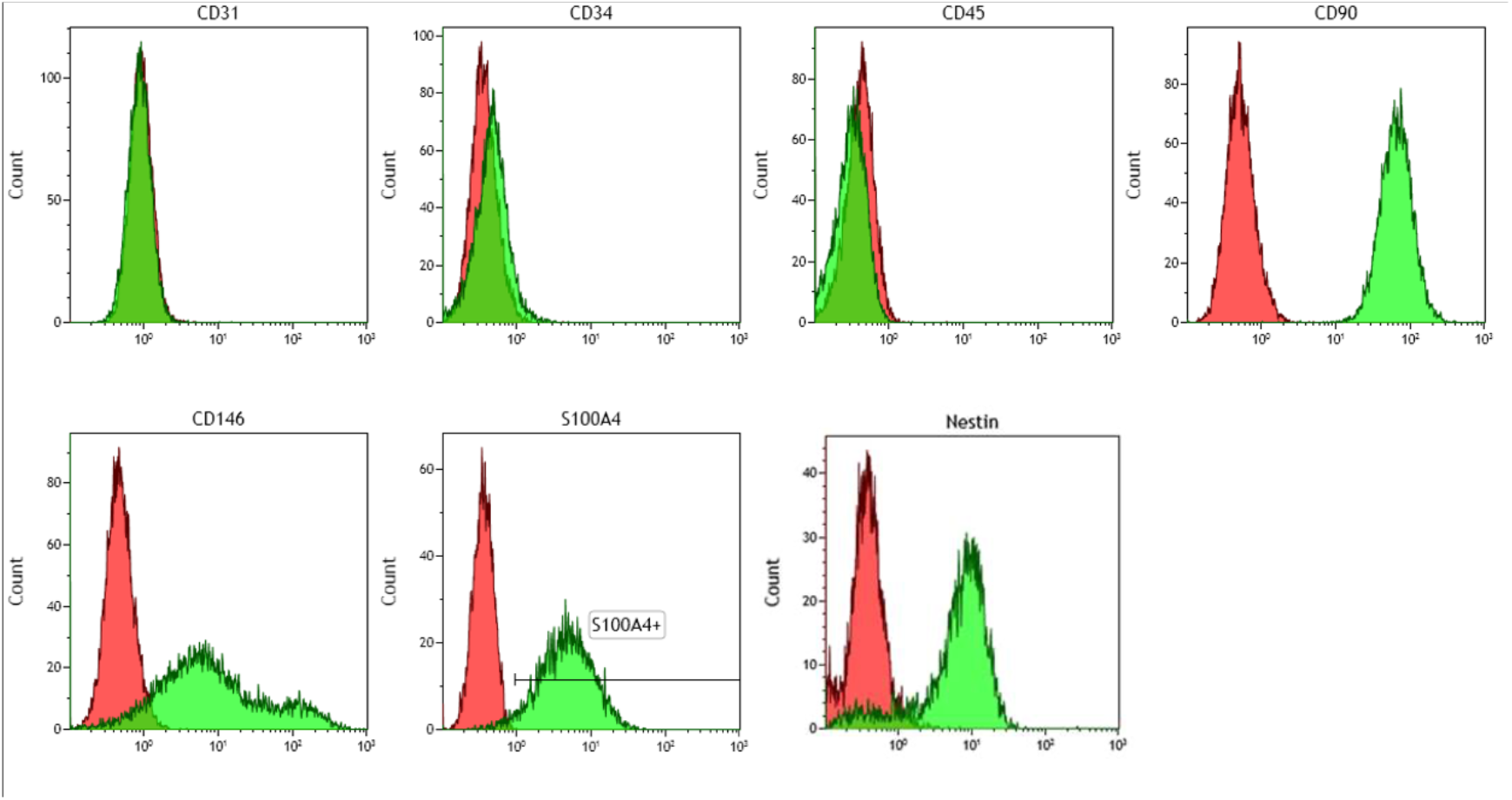
Phenotypic characterization of olfactory stem cells. Flow cytometry indicates that purified cells are negative for the usual markers of hematopoietic stem cells (CD31, CD34, CD45) and positive for the recognized markers of olfactory ecto-mesenchymal stem cells (CD90, CD146, S100A4, NESTIN).

### 3.2 Production of olfactory ecto-mesenchymal stem cells extracellular vesicles efficient for neurogenic enhancement

#### 3.2.1 Purified vesicles express the recognized markers of EVs

Figure 5A indicates that the number of EVs increases with time, reaching the total figure of 150 million at Day 4. In the meantime, vesicles from the culture medium strongly decreases. The mean diameter of secreted vesicles - 262±96.8 nm – confirms that the population is mostly composed of large vesicles and not exosomes (Figure 5B). Western blotting indicates that EVs express CD9, a specific vesicle marker. Figure 5C also shows that EVs are CD63^+^, INTEGRIN α5^+^, GAPDH^+^, CAVEOLIN^+^ and ALBUMIN^-^. Stem cells display a similar pattern of protein expression, except for albumin.

**Figure 5.**
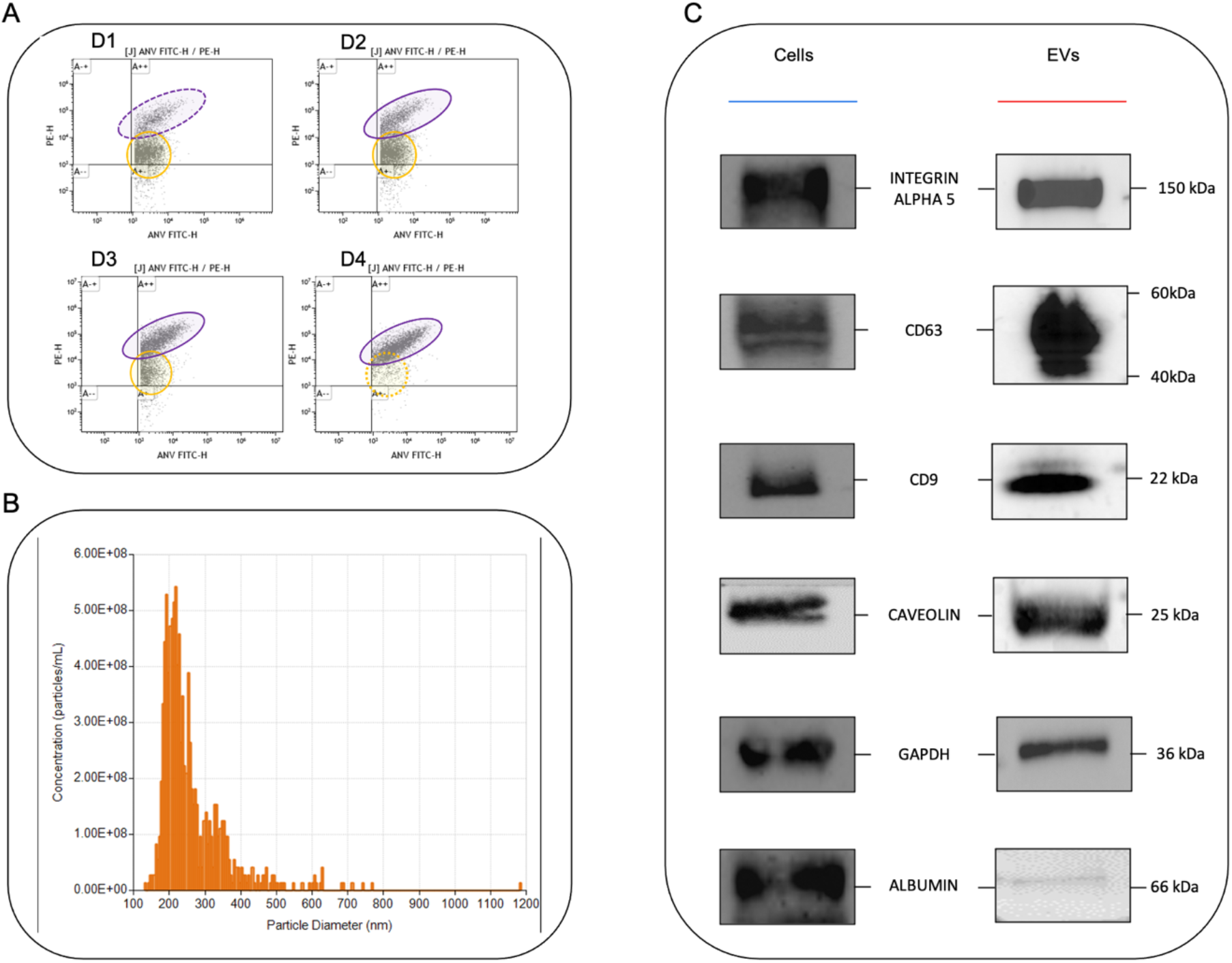
Characterization of OE-MSC-derived extracellular vesicles. A) Identification of extracellular vesicles (purple oval), at Days 1, 2, 3 and 4, using an anti-annexin 5 antibody. The number of EVs increases over time, reaching the number of 150 million. In the meantime, the number of contaminating vesicles (yellow circle), originating from the platelet lysate, decreases. B) Quantification and size estimation of EVs. The mean diameter of secreted vesicles is estimated to 262±96.8 nm. C) EVs express the recognised vesicle markers (CD9, CD63, INTEGRIN α5, GAPDH, CAVEOLIN) and are negative for albumin. Stem cells display a similar pattern of protein expression, except for albumin.

#### 3.2.2 Extracellular vesicles promote cell differentiation and neurite elongation

When incubated for several days with EVs, N2a cells tend to differentiate into neurons. When compared with cells cultivated with the differentiation medium, the ratio of differentiated cells is significantly increased at Day 4, at the following EVs concentrations: 200,000/3.6 cm^2^ well and 2×10^6^/3.6 cm^2^ well (Figure 6). In addition, EV-incubated cells extend longer axons at Day 4 compared to the differentiation medium, whatever the vesicle concentration used. This effect is noticeable as soon as Day 3, when EVs are present at the concentration of 20,000 (Figure 6).

**Figure 6.**
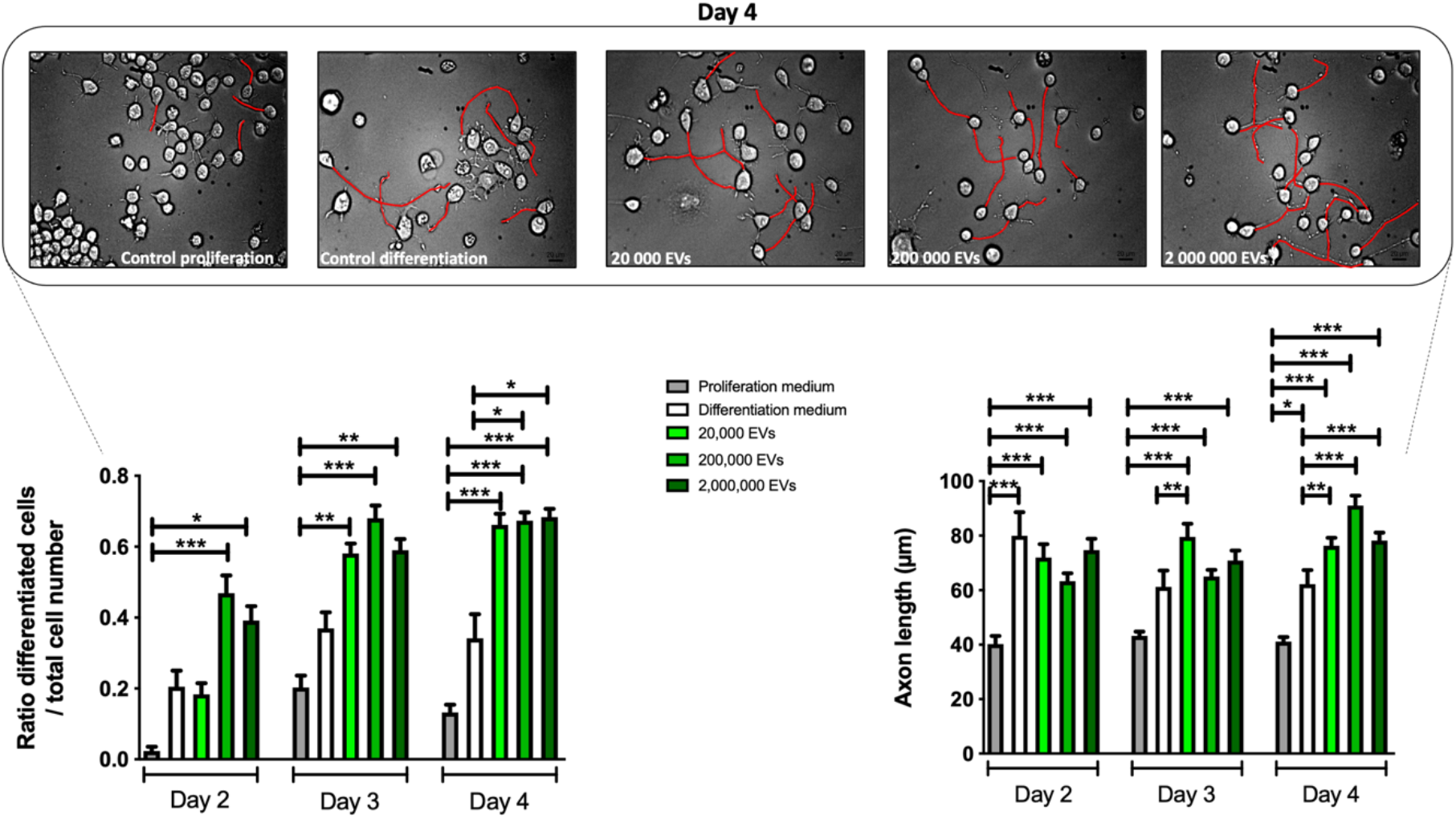
Effect of OE-MSC-derived extracellular vesicles on neuronal differentiation. The percentage of differentiated N2a is significantly increased at Day 4, when 200,000 or 2×10^6^ EVs are inserted into the culture medium. E) Similarly, axon elongation of N2a cells is significantly augmented at Day 4, whatever the vesicle concentration used. This effect is noticeable at Day 3, when EVs are present at the concentration of 20,000/ 3.6cm^2^ well.

## 4 Discussion

The current study reports a newly devised protocol for the purification and characterization of human OE-MSCs and their extracellular vesicles. The manufacturing process is in line with the rules and guidelines of national and European health agencies and the produced cells and vesicles can be considered as advanced therapy medicinal product (ATMP) that can be used in clinical trials. In addition, we demonstrate *in vitro* that OE-MSC-derived EVs are potent inducers of axogenesis.

### A safer and more efficient manufacturing process for clinical-grade olfactory stem cells

Clinical trials based on investigational medicinal products require a manufacturing that complies with “Guidelines on Good Manufacturing Practice specific to Advanced Therapy Medicinal Products”. These guidelines are edited by the French national agency for the safety of medicines and the European commission with the regulation n° 1394/2007. To fulfill these requirements, several modifications were applied. First, all cultures were performed in the cell therapy laboratory of the local public hospital (AP-HM) that got the authorization to produce ATMPs from the French national agency for the safety of medicines (EudraGMDP n° TIE/20/O/001). Recurrent visits by the agency’s managers made it possible to maintain this high level of requirement. Various steps of the culture protocol were modified to i) suppress the use of xenogeneic products, ii) simplify procedures and iii) ease the fulfillment of GMP requirements. For example, the usual bovine serum was replaced by human platelet lysate, a potent surrogate for cell proliferation (42). Likewise, for cell passaging, the recombinant enzyme TrypLE, efficient for dissociating other mesenchymal stem cells (62,63), was preferred to the usual porcine trypsin/EDTA (11,16) that cannot be used for clinical-grade purposes (45). We demonstrated that both enzymes gave the same results on the proliferation and viability of olfactory stem cells.

In addition, the protocol has been adjusted in order to respond to clinical demand. To avoid a prolonged delay that could adversely affect the therapeutic outcome (46), clinicians insisted upon obtaining the highest number of cells in the shortest time. We succeeded in this task by inserting an enzymatic dissociation phase and using very efficient agents for cell proliferation.

### Enzymatic isolation is more efficient to produce more rapidly human OE-MSCs

In most of our previous studies, human olfactory stem cells were purified by chopping the olfactory mucosa and placing each small explant under a glass coverslip. This technique was elected because the human olfactory lamina propria is thick and compact and all tested enzymes always led to an incomplete dissociation (7, 23). As a general rule, when compared to enzymatic methods, the explant technique leads to a better outcome with a reduced heterogeneity of cell populations, an increased cell proliferation and an improved cell viability. Unsurprisingly, the enzymatic isolation induces a stress for the cells while the explant protocol allows to preserve an extracellular matrix, a highly favorable substrate for cell survival and migration (47).

The explant technique is less time consuming and more cost effective (48). Even more so when the enzymatic dissociation requires the addition of expensive growth factors (FGF 2, PDGF…) in the culture medium. However, the purification of stem cells grown under coverslips requires additional GMP validations that, at this stage, remain uncertain.

For the first time, we used a health agency-approved enzyme and, according to the post-dissociation cell count, the result was similar to those obtained with other collagenases (Murrel et al, 2005, Matigian et al, 2010, di Trapani et al, 2013). We also compared explantation and dissociation techniques for the first time. We observed that although the enzymatic digestion was incomplete confluency was reached faster with the latter protocol. Such a result is the consequence of a double phenomenon: some time is necessary for the cells to migrate out of the explant while the dissociated cells can spread over the whole flask.

We also compared cell viability and phenotype, according to the two techniques of purification. As opposed to other MSCs (48), the enzymatic dissociation provides more viable OE-MSCs at day 21 (passage P2) and induces the same phenotype. In the current study as well as in a previous one (18) purified human OE-MSCs are CD146-positive, a marker of multipotency. Intriguingly, this cell surface marker was found not expressed in the seminal study dedicated to the characterization of OE-MSCs (11).

### Platelet lysate is a powerful enhancer of OE-MSC proliferation

Bovine serum is routinely used for MSC expansion but, when cell production in clinical GMP conditions is considered, the use of animal derivatives is discouraged as it can be a source of xenogeneic antigens and zoonotic infections (49–51). Platelet lysate is now a new standard for GMP-compliant cell manufacturing, particularly when serum-free fully defined media are not yet available for specific cell types (52). Here we demonstrate that platelet lysate improves the in vitro proliferation of olfactory stem cells. Previous studies report similar results for mesenchymal stem cells (BM-MSC, adipose derived MSC, umbilical cord MSC, dental pulp MSC, corneal stromal MSC) (53–60). In addition, it has been shown that platelet lysate maintains the differentiation potential, the immunomodulatory activity, and the telomere length and chromosomal activity of the mesenchymal stem cells (61). It also supports beneficial clinical outcomes (62).

The efficacy of PL is probably associated with its rich content in plasma-borne substances and specific platelet-derived growth factors, cytokines and chemokines (52). However, so far, no work focused on olfactory stem cells and the current study, mainly based on quantitative criteria, requires further investigation into the quality of the cells produced.

Previously, it has been demonstrated the optimal concentration of platelet lysate ranges from 5% to 10% when proliferation is considered and from 15% to 20% when osteogenesis and adipogenesis are assessed (58,63–65). Nevertheless, the results vary depending on the type of MSC and the mode of platelet lysate production. The content of commercial platelet lysates fluctuates according to the source material (fresh blood or not), preparation methods (repeated freeze/thawing cycles, direct platelet activation by calcium chloride, sonification or solvent/detergent treatment), sterilization procedure (irradiation, pathogen inactivation)… In the current study, platelet lysate PL100 contains 100% of plasma whereas PL30 contains only 30%, herein LP100 was more effective to stimulate cell proliferation than LP30. At least one study (66) also showed that PL supplemented with plasma is more efficient than unsupplemented PL in cell expansion. Like PL, plasma contains growth factors, therefore the percentage of plasma is an important factor.

Proven to effectively inactivate viruses (67), gamma irradiation does not affect the proliferation of OE-MSCs, as demonstrated here and in a previous study that assessed its role on stem cell proliferation (BM-MSC) and the content in growth factors as well as their clonogenic and differentiation potential and immunosuppressive properties (67).

### A process for manufacturing extracellular vesicles that meets very strict criteria/guidelines

Over the past decade, basic and clinical researchers focused their attention on the characterization of extracellular vesicles produced by stem cells. They also assessed their potential in diagnosis, monitoring, evaluation, prognosis, and therapy. Three teams previously isolated and identified extracellular vesicles from olfactory stem cells (34–36). In continuation of these works, this study describes (1) a purification method that meets the criteria required by EV specialists and European and French health authorities; (2) the effects of EVs on axogenesis of neural cells originating from neuroblastoma.

In 2018, the international society for extracellular vesicles (ISEV) published guidelines for studies on EVs (Minimal Information for Studies of Extracellular Vesicles (“MISEV”) guidelines)(68). As recommended, we measured the size, quantitated the numbers per million of cells and identified at least 3 positive and one negative markers for OE-MSC EVs. Our results are in line with previous studies (34–36), except for the expression of CD9 and CD63. Of note, the positive expression of these two markers is in accordance with one of our previous studies showing the production of these molecules at the transcript and protein levels (11).

### OE-MSC-derived extracellular vesicles promotes axogenesis

In two previous studies on rat models of peripheral nerve injury, we demonstrated the efficacy of OE-MSCs in improving motor functional recovery of the lower limb or face (21,22). To assess whether axonal regrowth was associated to extruded factors, a neuroblatoma cell line (N2a) was cultivated with a 50:50 mixture of fresh medium and the same medium collected from a 48 hour-long culture of OE-MSCs. We observed that the secretome of OE-MSCs induces an axonal elongation at least equal to that obtained with the standard differentiating medium (unpublished data). These promising results led us to move further and evaluate the role of the extracellular vesicles, today considered as the main actors in regenerative medicine (69). According to a review article, based on 206 preclinical studies on MSC-derived extracellular vesicles, beneficial effects were observed in 97% cases (70). However, only four studies satisfied all the quality requirements listed by the MISEV 2018.

When compared to the usual differentiating culture medium for N2a, extracellular vesicles induce an enhanced axogenesis. We observed an increased number of differentiated cells and an augmented axonal length. This double improvement can only be explained by the quality of the molecules released by the extracellular vesicles. One team characterized the content of OE-MSCs EVs and identified 304 proteins with key roles in cell differentiation, neural repair, angiogenesis, and inflammation (71). In order to move a step further and identify the molecules associated to nerve regeneration, we are currently comparing the mirnome, proteome and metabolome of EVs originating from various mother cells of multiple donors.

In addition, vesicles can be enriched with therapeutical molecules to improve their efficacy. Several studies describe new methods for adding these effectors and, according to the above mentioned review article, experiments using EVs enriched with therapeutical molecules demonstrated an improvement in the vast majority (71%) of cases (36). Vesicular engineering can also guide vesicles toward a target organ or area. (72). In rats, intraperitoneal injection allows efficient delivery to the target tissue (adipose tissue) whereas intravenous injection leads to accumulation of EVs in the spleen and liver (73). From a clinical point of view, intravenous administration is sought after but remains little used because of the captation of EVs by the lungs. To improve the targeting of intravenously delivered EVs, a more advanced knowledge on the mechanisms of EV fusion with the cell membrane is essential. Strategies aiming to integrate relevant receptors or ligands on the surface of EVs need to be evaluated (74).

### A ten to one EV/cell ratio may be optimal

Taking into account preclinical studies in rats, we evaluated a possible dose-reponse relationship between vesicle number and N2a cell differentiation. In contrast to previously published experiments (70), no obvious correlation between dose and axogenesis was observed, for either the number of differentiated cells or axon length. However, further analysis of differentiation and elongation kinetics indicates that 200,000 EVs condition (for 30,000 N2a cells) provides the best ratios with the most durable effect, as early as day 2. These data are consistent with other articles reporting that the most favorable EV/cell ratio is 10/1 (68,70).

For this first experiment we favored the use of freshly isolated vesicles because it has been shown that storage can alter the efficacy of EVs, especially when associated with enzymatic activity (75). Nevertheless, EV banking is necessary in clinical practice to provide vesicles during the acute phase of injury. This implies testing the efficacy of frozen and thawed OE-MSC-EVs. Furthermore, we only tested a single administration of vesicles. However, nerve regeneration is a long process and repeated administrations may be beneficial. For humans, recurrent ultrasound-guided injections of EVs tailored to each phase of nerve regeneration may be considered.

### Limits of the study

The cell model used in this study (N2a) is a mouse neuroblastoma cell line which has two limitations. It is a tumoral cell line derived from the central nervous system. Although it is a well-known model (with SH5SY) for the study of neuronal differentiation and axonal growth, the relevance of its use for the peripheral nervous system can be questioned. The cellular mechanisms involved in peripheral nerve repair are specific, notably the myelinization performed by Schwann cells.

On the second hand, these cells come from mice. If vesicles have fewer histocompatibility molecules and allow allografts or even xenografts, the results presented here may still be biased. Furthermore, while differentiation of OE-MSCs into Schwann cells is not essential for stem cell efficacy (76), their therapeutic effect might be partly enabled by Schwann cells present in the lesion area. This leads to the consideration a co-culture of Schwann cells and neuronal cells.

Finally, our experiments on the efficacy of EVs for neuronal differentiation were conducted with EVs extracted from the most cell-productive donor in a limited time. However, quality does not necessarily correlate with quantity, and high EV secretion may be the result of stress on the mother cell (68). Our future experiments should involve a comparison of cells from multiple donors.

## Abbreviations

BM-MSC: Bone marrow mesenchymal stem cell
ENT: Ear Nose Throat surgeon
EV: Extracellular vesicle
FBS: Fetal bovin serum
LCTC: Laboratory of Culture and Cell Therapy
MSC: Mesenchymal Stem Cell
OE-MSC: Olfactory ecto-mesenchymal stem cell
OE-MSC EV: Olfactory ecto-mesenchymal stem cell extracellular vesicle
PL: Platelet lysate

